# Performance Assessment of Variant Calling Pipelines using Human Whole Exome Sequencing and Simulated data

**DOI:** 10.1101/359109

**Authors:** Manojkumar Kumaran, Umadevi Subramanian, Bharanidharan Devarajan

## Abstract

The whole exome sequencing (WES) is a time-consuming technology in the identification of clinical variants and it demands the accurate variant caller tools. The currently available tools compromise accuracy in predicting the specific types of variants. Thus, it is important to find out the possible combination of best aligner-variant caller tools for detecting SNVs and InDels separately. Moreover, many important aspects of InDel detection are not overlooked while comparing the performance of tools. One such aspect is the detection of InDels with respect to base pair length. To assess the performance of variant (especially InDels) caller in combination with different aligners, 20 automated pipelines were developed and evaluated using gold reference variant dataset (NA12878) from Genome in a Bottle (GiaB) consortium of human whole exome sequencing. Additionally, the simulated exome data from two human reference genome sequences (GRCh37 and GRCh38) were used to compare the performance of the pipelines. By analyzing various performance metrices, we observed that BWA and Novoalign aligners performed better with DeepVariant and SAMtools callers for detecting SNVs, and with DeepVariant and GATK for Indels. Altogether, DeepVariant with BWA and Novoalign performed best. Further, we showed that merging the top performing pipelines improved the accurate variant call set. Collectively, this study would help the investigators to effectively improve the sensitivity and accuracy in detecting specific variants.

## Introduction

In the field of human genetics, Whole genome sequencing (WGS) explores clinical utility that provides the collection of individual’s genetic variation and etiology of the disease. Yet, the recent focus is on Whole exome sequencing (WES), which is being standard approach and is economic [1]. Though it covers only exonic regions (<2% of the whole genome), it produces the large quantity of data (raw reads) that requires a significant amount of bioinformatic analysis to produce biologically meaningful information [2]. In addition, the output must be accurate (detecting definite variants) and consistent in the identification of variants that account for the impact on phenotype.

The first obstacle to the accuracy in variant detection is the technical error by exome capturing kits while capturing the regions of our interest. It increases the possibility of missing some significant variants [3]. Secondly, the detection of variants through *in silico* methods play a vital role. Though plenty of tools are available [4–5], each performs best with the data obtained from particular sequencing platform. To mention, SAMtools is best for Ion Proton data [6] and GATK is for Illumina data [7]. As the different tools may produce a different list of variants for the same input, the efficiency of the variant callers is still not clear [8, 9]. Currently, no single tool is superior in detecting all the definite variants. However, on the other hand, applying multiple tools can result in higher misleading output [10]. It has also been reported that read aligners influence the accuracy of variant detection [9,11]. Thus, it is important to evaluate the optimal combination of aligners with the variant callers that may produce accurate variant calls including single nucleotide variants (SNVs) and small insertion and deletion (InDels).

Several benchmarking studies have been performed to evaluate the variant calling pipelines, which employs different aligners and variant callers. Notably, Liu et al. studied the performance of the pipelines with fixed aligner and different variant callers using single-sample and multi-sample variant-calling strategies. The study showed that GATK is found be a powerful variant calling method and SAMtools showed high true positive SNVs on simulated whole genome sequencing data [12]. In another study, the performance was analyzed based on read depth, allele balance and mapping quality using different variant callers, wherein GATK outperformed with low coverage data and yielded more accurate data for multiple-sample calling [13]. An in-depth comparison has been conducted only for SNV detection, specifically on somatic variation in cancer cells by Roberts et al. They have reported that the candidate SNV sets by different algorithms differ by number, the character of sites, sources of noise and sensitivity to low allelic fraction candidates [14].

Identification of InDels is more challenging because of the limitation of guidelines to detect them from sequencing data, compared to SNVs [15–16]. The most common InDel issues include low concordance rate among different sequencing platforms, realignment error, error near perfect repeat regions and incomplete reference genome in some cases [17]. Narzisi et al. suggested that the use of micro-assembly to reduce such errors [18]. Yet, the detection of large (basepair length) InDels are much difficult than small InDels [17,19]. Thus, it is essential to consider basepair length of InDels as an important parameter in variant detection.

In this study, we attempt to assess the best combination of alignment tool with variant callers for SNVs and InDels separately. To perform this, whole human exome sequencing dataset NA12878 from the public repository and the simulated data were taken as input. Five different aligners and 4 different variant callers were executed in all pairwise combinations (20 pipelines). Then we examined the variant calling capabilities of the pipelines by comparing their output with reported variants present in NA12878 exome. Majorly, this work aimed to build an extensive benchmark for studying the performance of pipelines in detecting SNVs and InDels.

## Results

In order to assess the performance of aligners and variant callers for detecting accurate SNVs and InDels from WES data, 20 pipelines were developed and the results were compared with gold standard variants dataset NA12878 provided by GiaB.

### Performance of variant callers

Initially, reads of human exome dataset NA12878 were checked for quality and adapter sequence was trimmed. Then, reads were aligned with the reference genomes GRCh37 and GRCh38 by different tools as given in Figure 1 (Details given in Table S1). After the post-alignment process, 4 different variant calling tools viz. GATK, SAMtools, FreeBayes and DeepVariant were used. Altogether using 20 different pipelines, SNVs were detected in 4 exome datasets (i) NA12878 aligned with GRCh38 genome (exome-1), (ii) simulated exome using GRCh38 genome (exome-2), (iii) NA12878 aligned with GRCh37 genome (exome-3), and (iv) simulated exome using GRCh37 genome (exome-4). The pipelines were run on our server (340GB RAM with 40 core for exome-1and -2; 320GB RAM with 32 core for exome-3 and -4) and the run time of each pipelines is displayed in Supplementary Table S2 for all 4 exomes. To access the performance of pipelines, we calculated true positive (TP), false positive (FP) and false negative (FN) variants using GiaB variant call set as standards. It contains 23,686 SNVs and 1,258 InDels for NA12878 exome. F-score was used as the function of performance.

**Figure 1.**
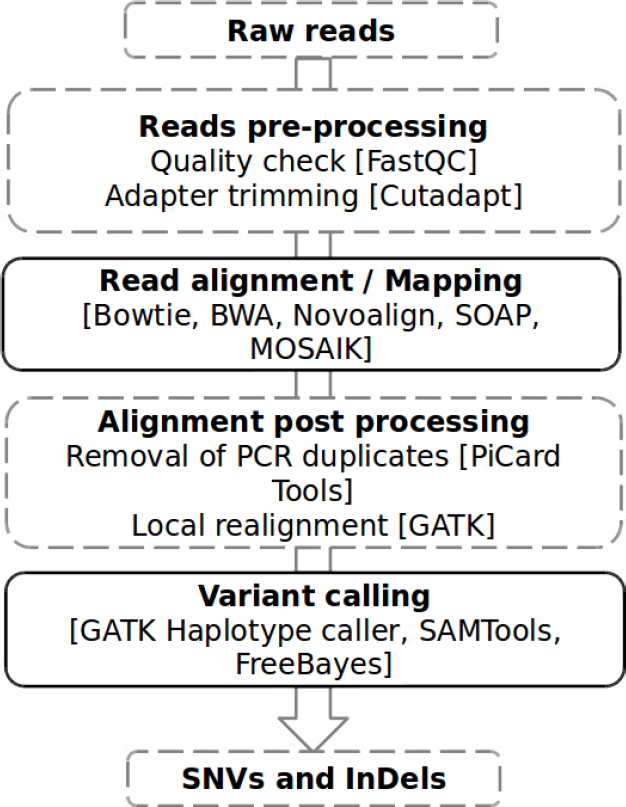
Schematic of the NGS data analysis pipeline.

In all the exome datasets, BWA_DeepVariant, Novoalign_DeepVariant, BWA_SAMTools, Novoalign_SAMTools were the top performing pipelines for the SNVs (Supplementary Table S3). The F-score of the top 4 pipelines were 0.97 on exome-1, 0.99 (except BWA_SAMTools) on exome-2, 0.98 (except BWA_SAMTools) on exome-3, 0.98 on exome-4. Next to these 4 pipelines, better results were obtained from Bowtie_DeepVariant, Bowtie_SAMTools and Mosaik_DeepVariant. In case of InDels, BWA_DeepVariant, Novoalign_DeepVariant scored best followed by BWA_GATK and Novoalign_GATK. Of these, DeepVariant based pipelines performed better than GATK based, showed the highest F-score of 0.99 on all exomes (Supplementary Table S3). Further, to explore how the sequencing depth affects the performance, the receiver operating characteristic (ROC) curves were plotted with the top 6 performing pipelines on all exomes. All the 6 pipelines performed roughly at the same level for SNVs, while subtle difference in their performance for InDels. Most of the SNVs and InDels were detected at about 150x, which indicated that this depth is a sufficient parameter for the variant detection analysis. However, Novoalign_SAMTools and BWA_SAMTools was less compared to others for InDels as expected (Figures 2 & S1).

**Figure 2.**
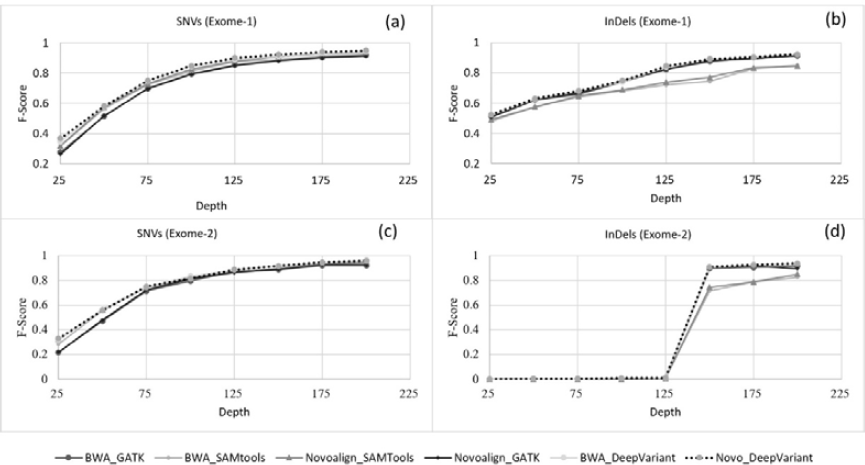
F-score as a function of depth for top 6 pipelines. ROC curves were plotted using the depth of SNVs (a, c) and InDels (b, d) against F-score using exome-1 (a, b) and exome-2 (c, d).

Next, we examined the F-score of each pipeline with respect to genotype quality (GQ) as a scale of variant detection performance. At GQ>60, the pipelines showed good performance for both SNVs and InDels on all 4 exomes (Supplementary Table S4). Of the top six pipelines mentioned earlier, BWA_DeepVariant and Novoalign_DeepVariant remains on top for both SNVs and InDels on all exomes (Figures 3 and S2). BWA_SAMTools and Novoalign_SAMTools followed next for SNVs and Novoalign_GATK and BWA_GATK for InDels. These pipelines increase their performance as GQ increases, suggesting that higher GQ increases the rate of performance with respect to F-score. Surprisingly, in InDels detection BWA_DeepVariant and Novoalign_DeepVariant showed increased performance even at GQ>60 on simulated exome data (Figures 3d and S2).

**Figure 3.**
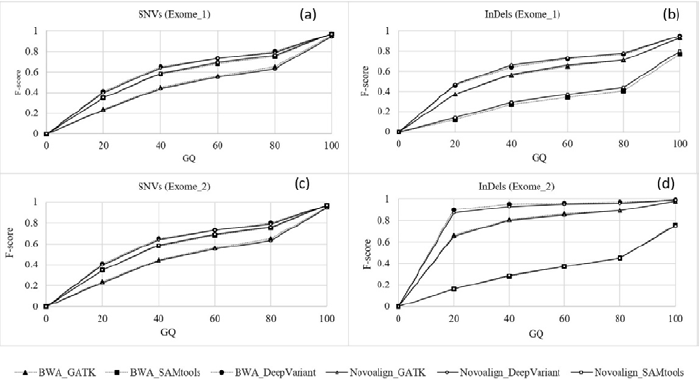
F-score as a function of genotype quality for top 6 pipelines. ROC curves were plotted using the GQ of SNVs (a, c) and Indels (b, d) against F-score using exome-1 (a, b) and exome-2 (c, d).

Further, we evaluated the accuracy of the SNVs and InDels with respect to genotype concordance. As expected, there was no change on top performing pipelines by comparing F-score as a function of genotype concordance on all exomes (Supplementary Table S5). We also investigated the ratio of heterozygous to homozygous (het/hom) and found the ratio was higher in SNVs detection than InDels. The ratio was ~1.6 for exome-1 and -2, and ~1.5 for exome-3 and -4 for SNVs. While ~1.2 for exome-1 and -2, and ~1.2 and ~1.3 for exome-3 and exome-4 respectively. Difference in the performance was observed when we compared the heterozygous and homozygous detection with respect to F-score (Supplementary Table S6). BWA_DeepVariant, Novoalign_DeepVariant, BWA_SAMTools and Novoalign_SAMTools showed high F-score, >0.96, for SNVs on all exomes. While BWA_DeepVariant, Novoalign_DeepVariant, BWA_GATK and Novoalign_GATK score high, >0.9, for InDels, and rest of them were failed to follow including BWA_SAMTools and Novoalign_SAMTools.

### Performance in SNV calling using Ti/Tv ratio

One of the key quality metrics of SNV call set is the ratio of transition (Ti) to transversion (Tv). The Ti/Tv ratio was ~3.4 for exome-1 and -2, while ~3.2 for exome-3 and -4. We also investigated F-score with respect to transition (Ti) and transversion (Tv) compared to gold standards. The pipelines Novoalign_DeepVariant and BWA_DeepVariant scored high for Ti and Tv on all exomes followed by Novoalign_SAMTools and BWA_SAMTools (Supplementary Table S7).

### Performance in InDel calling at different base pair (bp) length

We analyzed the InDel detection performance of the pipelines with respect to F-score as a function of insertion and deletion base pair length. DeepVariant and GATK pipelines, particularly along with the aligners BWA and Novoalign, showed increased InDel detection rate at higher base pair length on all exomes. However, the performance of pipelines were differed at particular bp length of InDels. In all the 4 exomes, majority of the tools found deletions at 17, 23, 25 and 26 bp and insertions at 22 and 35 bp length with F-score almost one (Figures 4, 5 and Supplementary Figures S3 & S4). BWA_DeepVariant and Novoalign_DeepVariant detected more number of insertions on exome-2 and -4. The deletions of 24 and 27 bp length were not detected by any of the pipelines on exomes-1, -3 and -4. While, insertions of 13, 23, 24, 27 and 59 bp length were detected by none on exomes-1 and -3. Also, the large 59 bp insertion was not detected on exome-2 (Supplementary Table S8). We pointed out the possible reasons for the InDels that is being not detected by the pipelines in the discussion section.

**Figure 4.**
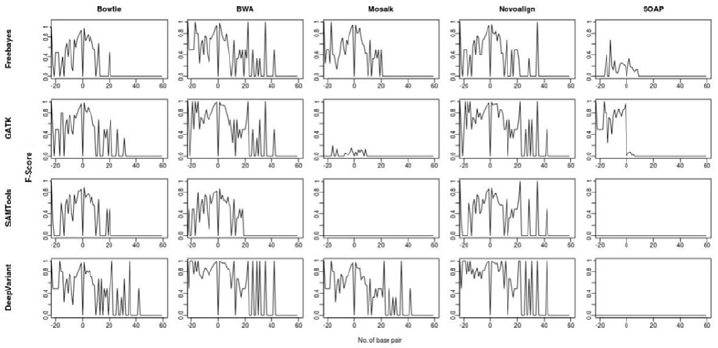
InDels detection performance in exome-1. F-score of InDels were plotted against the base pair length of the InDels. Negative value of x-axis indicates the deletion and positive value for insertion.

**Figure 5.**
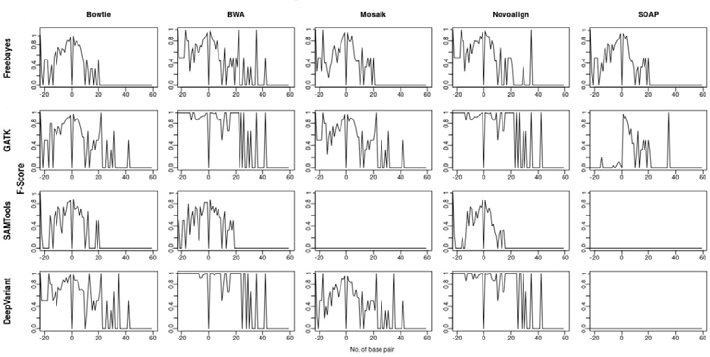
InDels detection performance in exome-2. F-score of InDels were plotted against the base pair length of the InDels. Negative value of x-axis indicates the deletion and positive value for insertion.

### Comparison of best performing pipelines

In order to improve the accuracy in variant detection, we compared the GiaB true call set against the variants detected by the top 4 pipelines (mentioned earlier) BWA_DeepVariant, Novoalign_DeepVariant, BWA_SAMtools, and Novoalign_SAMtools, for SNVs (Figure 6a, 6c) and BWA_GATK, Novoalign_GATK, BW A_DeepVariant, and Novoalign_DeepVariant for In Dels (Figure 6b, 6d). The similar were analyzed on exome-3&-4 and illustrated in Supplementary Figure S5.

**Figure 6.**
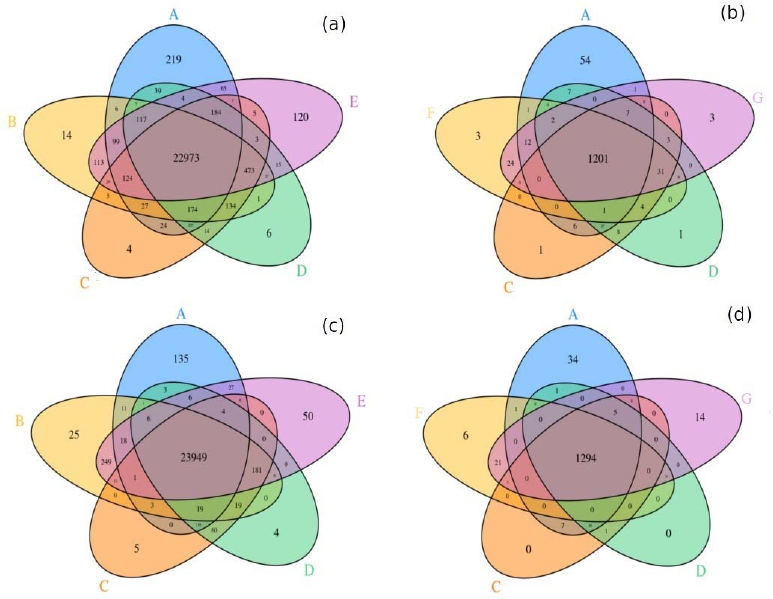
Venn diagram depicting the comparison of GiaB variants (A) against the top performing pipelines B. BWA_SAMtools, C. BWA_DeepVariant, D. Novoalign_DeepVariant, E. Novoalign_SAMtools, F. BWA_GATK and G. Novoalign_GATK for SNVs (a, c) and InDels (b, d) on exome-1 (top row) and exome-2 (bottom row).

The concordance of variants with GiaB call set was improved for all 4 merged pipelines on all exome (Figures 6 and S5). The accuracy in calling true positive SNVs were improved to ~99 % on exome-1 and -2, and ~98 % on exome-3 and -4. In case of InDels, ~96 % was observed on exome-1 and -3; however, ~98 % was observed on simulated exomes (exome-2 and -4). Further, we evaluated best performing caller DeepVariant by merging its call set from BWA and Novoalign alignments. This showed ~98% and ~96% TP for SNVs and InDels on all exomes respectively. Altogether, merged pipelines showed increased performance by improving the accurate variant call set, despite of the increased the FDR.

Although each caller uses different algorithms (strategy to identify the variants as given in Supplementary table S1), ~0.5-1.5% and ~0.5-4% false negative (FNs) SNVs and InDels were observed in all the exome dataset respectively. To investigate further, the genotype quality (GQ) and depth of the FN variants not detected by the top pipelines were analyzed. The depth of the FNs obtained by BWA and Novoalign alignments were plotted (Figure 7a-h) and the graphs showed they fell under the upper limit of 30X on all exomes. However, there are outlier suggesting that the depth might not be the only reason that affect the performance of variant detection. By comparing the GQ (Figure 7i-p), which showed <10 for all FNs on all exomes, the variant callers possibly missed the true variants due to the low GQ.

**Figure 7.**
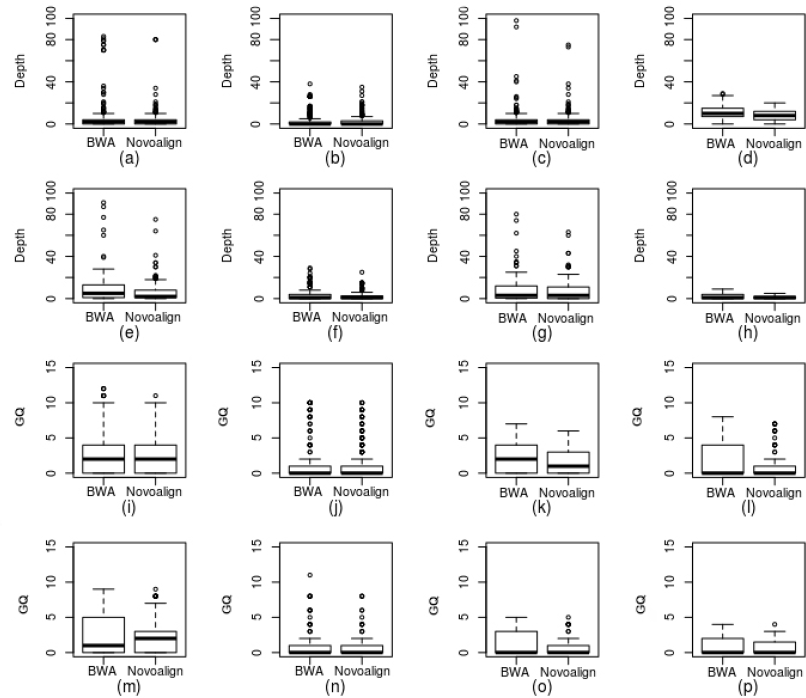
Analysis of depth and genotype quality of true SNVs missed (false negatives) by BWA and Novoalign alignments. Depth of the false negative SNVs on exome-1(a), -2 (b), -3(c) and -4 (d) and InDels on exome-1(e), -2(f), -3(g) and -4(h). Genotype quality of false negative SNVs on exome-1 to -2 (i, j, k and l) and InDels on exome-1 to -4 (m, n, o and p) respectively.

## Discussion

A major challenge in whole exome sequencing is how to process the data to yield high quality disease variants for downstream analysis. Currently, a variety of tools are available and we have automated these tools (differs in aligner and variant caller) to identify the best pipeline in variant detection. The human exome NA12878 was used for assessing the performance of the pipelines. Simulated data were also used, which is most popular for assessing and validating biological models and for understanding of specific data sets [20]. Even though there are differences among the results of these datasets, all the experiments are important as they give a different perspective when comparing the pipelines.

The overall performance ranking of the pipelines is similar in real and simulated data; however, the false discovery rate seems much lesser in simulated data, this might be due to the underlying error model of experimental exome data. The pipelines with BWA and Novoalign (non-commercial version) aligner members were attributed much in variant detection. The algorithm of BWA balances between running time, memory usage and accuracy. Whereas Novoalign is slow and high memory usage that contribute to better mapping. But results still varied greatly depending on the variant caller with respect to the specific variant which was called by the user.

In SNVs detection, Ti/Tv ratio was one of the metrics for performance comparison. The ratio was transiently in accordance to the reported ratio range of 2.6-3.3 [21], on exome-1&-2 and transiently accordance on exome-3&-4 except in SOAP_GATK on exome-3 (3.55) and in Mosaik_GATK on exome-4 (3.94). But the ratio may not always necessarily mean more accurate since low-frequency variants sometimes have higher Ti/Tv ratio than moderate frequency SNVs [7]. Also, as per principle proposed by McKenna et al. [22], in this study the more likely true variant set had the highest Ti/Tv value Thus, Ti/Tv ratio was taken for the accuracy of the true positive variant detection, and the overall performance was evaluated by F-score. It should be emphasized that in SNVs detection, the better performance was observed in GATK compared with SAMtools which was reported as the best performer earlier [12–13], but the present study reports SAMtools performs better than GATK for SNVs. However, the recently developed variant caller DeepVariant performs well compared to all other tools.

The overall performance of all the pipelines in InDel detection is comparatively lower than SNVs detection. This might be the result of using WES data as they miss many large InDels, and the accuracy of InDels detection with WES data is much lesser than that of WGS [17]. In the present work, in InDels detection DeepVariant member pipelines performed well compared to others both in accuracy as well as in higher coverage of base pair length. In respect to the aligner, BWA and Novoalign performed well in combination with DeepVariant. The investigation about missing variants reveals that genotype quality and depth may the reason for it, and the quality control process did not affect the detection of variants (Figure 7). In addition to it, in case of simulated exomes the error quality rate was set to 0.01% during simulation.

Taken together, DeepVariant can be used with BWA and Novoalign to achieve the more accurate call set. However, our recommendation is the use BWA and Novoalign aligners with DeepVariant and SAMtools callers for detecting SNVs, and with DeepVariant and GATK for Indels. However, the users may need to be aware that ~1% and ~2% of variants might not be detected of these pipelines. We conclude that our study will help to achieve more accurate variant calls, ultimately leading to the identification of disease causing variant for clinical genomics.

## Materials and methods

### Datasets

FASTQ files of human exome HapMap/1000 CEU female NA12878 (accession No.: SRR098401) was downloaded from NCBI-Sequence Read Archive (SRA- http://www.ncbi.nlm.nih.gov/sra). This was sequenced using HiSeq Illumina 2000 platform and SureSelect human all exon v2 target capture kit [23]. The target region BED file was downloaded from Agilent SureDesgin (http://earray.chem/agilent.com/suredesign, ELID: S0293689). The human reference genomes GRCh37 and GRCh38 were downloaded from the Ensembl [24]. GiaB high confidence callset version 2.19 along with a BED file was downloaded from NCBI, which was further filtered to highly accurate call set using the BED file. The list of variants provided in GiaB was created by integrating 14 different datasets from five different sequencers, and was used as ‘gold standard’ to validate the variants detected by pipelines.

Further to test the certainty of the performance of the pipelines, we have also used simulated human whole exome data generated by ART toolkit [25]. ART takes a reference genome in FASTA format and generates ‘synthetic’ sequencing reads. This mimics the technology-specific sequencing process with customized read length and error characteristics. The reference genomes GRCh37, GRCh38 and sequencing target BED (SureSelect human all exon v2 target capture region) file were inputs of the simulator. We have generated simulated short paired-end reads of 150 bp length with the depth of 150X covering sequencing targets for Illumina HiSeq 2000 sequencing technology with 0.01 % error model.

### Pipeline development

The modular pipeline (Figure 1) was developed to process both the original NA12878 data which were aligned with the genomes GRCh37, GRCh38 and their corresponding simulated exomes. The pipeline involves several steps to produce high-quality alignment files, and to predict definite variants. Initially, the quality of the raw reads obtained from SRA was checked by FastQC [26] and the low-quality reads, adapter contaminates were trimmed by Cutadapt [27]. After aligning the reads with the reference genome, PCR duplicates were removed using PiCard Tools [28]. To avoid pseudo SNVs and InDels, local realignment was done by GATK. This was followed by read recalibration, based on target capture region provided by Agilent Sure Design, for error detection towards obtaining the base quality. Finally, SNVs and InDels were identified by different variant calling tools. For alignment (read mapping) and variant calling, respectively 5 and 4 different tools (Supplementary Table S1) were used based on prevalence and popularity. All the steps were integrated using shell scripts (available from https://github.com/bharani-lab/WES-pipelines.git) and all the possible 20 pipelines were developed with default parameters.

### Performance evaluation of variant callers

The variants determined by pipelines were compared with standard variants provided by GiaB using VCFTools [29]. The performance of variant detection by different pipelines was measured statistically as, Sensitivity = TP / (TP+FN), Precision = TP / (TP+FP), False discovery rate (FDR) = FP / (TP+FP) and F-Score = 2TP / (2TP+FP+FN) where, TP is true positive variant found in both GiaB validated dataset and data determined by pipeline; FP is false positive variant determined by pipeline but not validated by GiaB; FN is false negative variant, known as missing variant which is validated by GiaB but not determined by pipelines. To analyze the performance, we used different metrics including read depth, genotype quality, genotype concordance and Het/Hom ratio. Furthermore, Ti/Tv ratio was used for SNVs and F-score as a function of base pair length for InDels. Venn diagram was plotted to compare the performance of top performing pipelines.

## Acknowledgment

The authors acknowledge the financial support of this work for Science and Engineering Research Board, Govt. of India (SB/YS/LS-97/2014).

